# Astrocyte-like CEPsh glia respond to stress

**DOI:** 10.1101/2022.11.03.515075

**Authors:** Evangeline Chien, Angel J. Garcia, Noelle D. L’Etoile, Rashmi Chandra

## Abstract

While astrocytes are known to be important for development and nourishment of the nervous system, the field is just beginning to explore how astrocytes respond to environmental stimuli. Using *Caenorhabditis elegans* and their astrocyte-equivalent, CEPsh, we asked whether astrocyte-like glia respond to odor exposure. We found one day-old adult *C. elegans* decrease *hlh-17* promoter (CEPsh glia marker) mediated fluorescent expression when trained with an innately attractive odor butanone. Moreover, the olfactory training paradigm itself affects p*hlh-17* expression, but in a different way. This suggests astrocyte-like CEPsh glia can integrate environmental information to respond to changes in the environment, which enhances survival.

## Description

Astrocytes are a type of glia that play a key role in numerous functions of the central nervous system (Sofronview and Vinters, 2010). Recently, astrocytes were also shown to participate in spatial memory formation in mice (Hosli et al,. 2022) and may respond to interior information such as dopamine (Pittolo et al., 2022). The nematode *Caenorhabditis elegans (C. elegans)* is a good model organism for such studies; it is transparent, allowing live imaging, and its CEPsh cells are well-described astrocyte-like glia (Shaham, 2015). Though the *C. elegans’* sensory ensheathing glia (AMsh) have been shown to detect odorants and drive olfactory adaptation (Duan et al., 2020), it remains unknown if their astrocytes respond directly or indirectly to environmental stimuli. Therefore, we asked whether the CEPsh cells are modulated during a spaced the olfactory training paradigm in which animals learn to avoid the innately attractive odorant, butanone by repeatedly pairing the odor with starvation and exercise (Chandra et al., 2022; Kauffamnn et al., 2011). The memory from this training remains for at least 24 hours (Chandra et al., 2022). In particular, we asked whether the cephalic sheath (CEPsh) glia, the astrocyte equivalent in *C. elegans*, integrate information from outside the animal with the animal’s internal, physiological, state. Our novel finding is that CEPsh glia do respond to odor-training by down-regulating expression of a CEPsh-specific gene. Thus, the astrocyte-like glia of *C. elegans* have the capacity to integrate external and internal signals which could lead to durable changes in the animal’s response to an external stimulus as a result of its experience with that stimulus.

p*hlh-17* was shown to be a CEPsh specific promoter (Rapti et al., 2017) whose expression we found, surprisingly, changes in our training paradigm (panel B training and D, E expression). We used the construct in Panel A, which we engineered to express the miniSOG and wrmScarlet operon under the control of the 2556 bp of sequence upstream from *hlh-17* start site (McMiller and Johnson 2005). In order to monitor the CEPsh cell’s response to the stimuli of training (swimming-exercise and starvation in buffer-trained cohorts, and swimming-exercise, starvation, and odor in odor-trained cohorts) we used a strain that carried an extrachromosomal array containing this construct as our reporter transgene for transcription in the CEPsh (Panel A).

Butanone is normally an odor that is associated with food (Worthy et al., 2018), sensed through AWC sensory neurons (reference), and *C. elegans* are innately attracted to it (Bargmann et al., 1993). Over three training cycles (T1, T2, T3) in which the worms were exposed to odor diluted into buffer without food, this attraction is converted to repulsion. We trained a control group of worms in the same way except that Butanone was left out of the Buffer solution over the same three training cycles (Panel B).

Glia ablation interfered with AWC-mediated chemotaxis (Bacaj et al., 2008), so it was important to test whether the reporter construct affected CEPsh function. We asked whether the reporter array affected the animals’ behavior by comparing the reporter expressing strains’ ability to learn and remember like wild type (N2) parents. In order to understand if any of our observations were due to off-target effects from our reporter transgenic array, we asked if wildtype, non-transgenic, and transgenic worms behave similarly. After the training sequence was complete, each group’s ability to learn and remember that butanone is not profitable was tested with the Chemotaxis Assay (Panel B Test Learning and Test Memory). We used the Chemotaxis Assay twice, once directly after T3 was completed (t=0) which tests learning and again after allowing the worms to rest and feed for 16 hours (t=16) which tests for long lasting memory. Each group was put onto a plate (panel B Test Learning) with two odors: diluted butanone and ethanol (a neutral odor). Sodium azide (a paralytic agent) was added to each spot on the assay plate. The worms ran for at least two hours before we counted the number of worms at each odor. The buffer-trained worms (control group) collected at butanone reflecting their attraction to this odor. The butanone-trained worms (case group) collected at buffer which reflects that the population has learned to avoid butanone (panel C Chemotaxis Graph, see below the graph in C for the calculation of the Chemotaxis Index which reflects the attractiveness of butanone). There was no significant difference detected between wildtype and transgenic and wildtype and non-transgenic across every condition and time point (see panel C). That is, neither learning (t=0) nor memory (t=16) was affected in the line that expressed the CEPsh reporter. Thus, this reporter strain behaves like wildtype as shown through chemotaxis assays, and/or odor-dependent learning.

In order to measure the expression of p*hlh-17* after each training cycle (T1, T2, T3), we imaged the fluorescence intensity Fluorescence in butanone-trained worms (panels D and E). We found that the reporter intensity in the CEPsh decreased compared to naive worms after the first training (T1) (*** p = 0.0006, Naive to Butanone T1). By contrast, fluorescence in buffer-trained worms did not significantly decrease until after the second training (T2) (** p=0.0018, Buffer T1 to Buffer T2). In each case, after the drop, fluorescence levels remained depressed after the subsequent cycles of training (ns between Butanone T1, T2 and T3; ns between Buffer T2 and T3). Additionally, after the drop, the fluorescence levels *between* the conditions were not significantly different at the comparable timepoints (ns between Buffer T2 and Butanone T2; ns between Buffer T3 and Butanone T3). Thus, buffer-trained and butanone-trained each showed a significant sustained reduction in fluorescence intensity, but the reduction was observed at different time points, with a faster response in the butanone-trained groups.

The significant drop in fluorescence intensity after training demonstrates that CEPsh glia respond to changes in the worm’s environment and possibly also respond to the worm’s internal state by decreasing expression from p*hlh-17*. The presence of the external stimulus, butanone, caused the CEPsh to respond more rapidly than when butanone was left out of the training (buffer). This means that they could be integrating external stimuli (odor) with their internal state (such as energy depletion and starvation). If this integrated response is transmitted to the neurons of the circuit, it may be causing change in the animal’s behavior.

### Future Directions

Given that we have now shown that the astrocyte-like CEP sheath glia responds to butanone in the environment, we can now explore how CEPsh receive information. Three of the possible ways CEPsh may receive information from the environment include: from neurons, specifically the butanone sensing AWC neurons; from AMPsh glia; and or, they may receive information directly from the environment. The astrocyte-like CEPsh may also sense changes internal to the animal that result from either starvation or exercise or both. Future experiments could identify whether starvation or exercise or their combination affects CEPsh gene expression. Additionally, we will ask how odor exposure might influence these internal states. To assure ourselves that the reporter does not produce artificial responses due to the expression of the blue light-inducible superoxide producing miniSOG, we quantified wormScarlet expression. To further ratify these results, we plan on deleting the miniSOG from our reporter construct and repeat these experiments, and/or use a different phlh-17:GFP construct. Insertion of a fluorescent reporter into the *hlh-17* gene itself using a CRISPR strategy would also clarify whether the extrachromosomal array affects the readout. Furthermore, other CEPsh expressed genes, in addition to hlh-17, should be looked at to see how they might be regulated by training. hlh-17 is likely not unique, and many genes may change expression in the CEPsh glia when the animals undergo the training paradigm. Understanding what these genes encode for and what they do could provide insight into what processes the training induces within the astrocytes. Nonetheless, we have identified a paradigm in which the CEPsh astrocyte-like glia respond to both an internal state change and external stimulus and this may indicate a previously unexplored capability of astrocytes to integrate stimuli.

## Panel Descriptions

A. Schematic representation of the gene construct that is driving miniSOG and wormScarlet expression through the *hlh-17* promoter, and the populations assayed throughout this study.
B. Olfactory training paradigm and measurement of fluorescent intensities. One group of worms were trained to associate Butanone (BTN) with starvation and exercise over three training sessions (T1, T2, T3). Second group of worms were trained in a neutral Buffer solution (Buff) over the same three training sessions. Between the three trainings, we provided the worms in each group with food (F1, F2). After each training (T1, T2, T3), we imaged the fluorescence intensity of individual worms to measure the expression of p*hlh-17*. After training, we tested each group’s learning on chemotaxis plates using BTN (odor) and EtOH (control). We counted the number of worms that are attracted by BTN. We allowed the worms to rest and feed for 16 hours and then tested their memory. The equation for measuring fluorescence in each animal is under the example image.
C. Chemotaxis Index (CI) measures how strongly worms move towards the attractive butanone odor. The equation for calculating CI is under the graph. The One-way Anova showed the transgenic strain behaved similarly to wild type animals.
D. Graph of fluorescence levels after each training cycle. After T1, the fluorescence of butanone-trained worms significantly decreased compared to naïve worms. By contrast, the buffer-trained worms showed no significant difference in fluorescence intensity until after T2. The table below the fluorescence graph lists the p-values for the statistically significant differences.
E. Representative images of transgenic *C. elegans* brain with fluorescent CEPsh glia for all cases: naive, Buff T1, T2, T3, and BTN T1, T2, T3. White line on composite images represents 50 microns.
F. Perspective: We found an astrocyte-like glia in C. elegans uses a specific 2.55 kb upstream element of hlh-17 gene to respond to training paradigm and odor addition differently. This raises the possibility that CEPsh glia may form olfactory memory through specific changes in gene expression that happens during integration of different environmental cues.

## Methods

### C. elegans strains and maintenance

N2 (wildtype) and an extrachromosomal array (JZ3003, p*hlh-17*::miniSOG::SL2::wScarlet at 50ng/uL and unc-122p::mcherry at 25ng/uL), were propagated on separate nematode growth media (NGM) plates under standard conditions. The plates were seeded with OP50, an *E. coli* strain. The plates were kept at room temperature, 20°C. Worm colonies were chunked or picked onto new plates every two to four days to prevent starvation (Wormbook protocols).

### Food for Long Term Memory (LTM) Assays

The day before each assay, we inoculated a single colony of bacteria in 100 mL of LB in an Erlenmeyer flask, leaving it for 16 hours in the shaker at 37ºC. The day of the assay, we poured 50 mL of the OP50 LB into a conical tube and centrifuged at 4000 rpm at 20 ºC for 20 minutes. We then resuspended the pellet in 16 mL of S. Basal and mixed until homogenized.

### LTM Assays

3-4 larval stage 4 (L4) worms were picked onto 2-4 agarose plates with NGM and an OP50 lawn per strain were picked 5 days in advance of the day of the Long-Term Memory (LTM) assay. On the day of the LTM, we screened the plates for any mold or bacterial contamination. We washed each plate with 1-2 mL of S. Basal into four 1.5 mL microcentrifuge tubes. After washing the worms three times, we added 500 μL butanone solution (5.5 μL 2-butanone and 50 mL S. Basal) to the butanone-trained tubes and added 500 μL S. Basal to the buffer-trained tubes. Then, the tubes were put on a rotisserie for 80 minutes. After 80 minutes, we washed the worms three times, added 500 μL of food, and put them on the rotisserie for 30 minutes. We repeated the 80 minute training three times interspaced with food for 30 minutes.

### Odor Attraction Chemotaxis Assays

We first poured the chemotaxis plates by boiling 1.6 g of bactoagar (Fisher Scientific, Cat # DF0140-01-0) in 100 mL ddH_2_O until no visible granules remained. Once it cooled to 55°C, we added 500 μL 1M K_3_PO_4_ (Sigma-Aldrich Cat # P37786 K_2_HPO_4_ and Cat # P5655 for KH_2_PO_4_), 100 μL 1M CaCl_2_ (Sigma-Aldrich Cat # C4901), and 100 μL 1M MgSO_4_ (Sigma-Aldrich Cat # M7506) and then pipetted 10mL of the gel per 10 cm petri dish. 15 minutes after pouring the gel into the dishes, one for each condition, we drew the origin and two choice boxes on the back of each dish (as shown in Panel B). 1 μL of 1M sodium azide (Millipore Sigma Cat # 2002) was added to the center of each box. 2-butanone was diluted 1:1,000 in 100% 200-proof ethanol. Right before plating the worms, 1 μL 100% ethanol and 1 μL butanone odor were added to their respective boxes. The worms were then washed with S. Basal three times and then plated at the south end of the plate. Worms were gently dabbed dry with a Kimtech wipe. The plates were used within 3 hours of being poured. We plated at t=0 and at t=16 hours after T3 to test learning and memory respectively. When the worms stopped moving, we counted how many were in each choice box and how many were outside.

### Fluorescence Quantification

We took images on a spinning disk confocal scope. After each 80-minute training had ended, we took a sample (∼25-50 worms) and placed them on an agarose pad with 10 mM tetramizol hydrochloride (Sigma Cat # L9756). We imaged using the Wittman 40x water lens (NA 0.75) and the Nikon NIS Elements Analysis platform. We imaged using the 561nm filter and the 488nm filter at an exposure of 100 ms with 25% laser power. To image the worms, we focused on the head of transgenic worms, found the upper and lower boundaries of the worm and created a z-stack of images, step-size of 0.3 microns. After the image was taken, we quantified the fluorescence level of the CEPsh glia using the Nikon NIS Elements Analysis platform. Specifically, we manually scrolled through the z-stack, selecting the slice where the greatest intensity was recorded according to the LUT intensity scale. For this slice of greatest intensity, a region of interest (ROI) was drawn around the ventral glia, and NIS Elements quantified the intensity. That same ROI was duplicated and moved to a place on the worm that had no added fluorescence other than auto-fluorescence. The fluorescence of the background ROI was then subtracted from the fluorescence of ROI_glia_ in order to get the net fluorescence of the glia. This process was repeated for the dorsal glia. The average was taken of the two ROI_glia_ in order to get the glia fluorescence level of the worm. If fluorescence was only detected for one glia, the second ROI_glia_ was set to be 0.

### Statistical Analyses

Statistics were performed using GraphPad Prism 9. P values used for the statistical readouts are ns is P>0.05, *P<0.05, **P<0.01, ***P<0.001, and ****P<0.0001. Each independent trial/assay was performed on a population of >50 worms that were picked and raised on different growth plates.

For the Odor Attraction Chemotaxis Assay, a Tukey One-way Anova was performed to statistically qualify that the transgenic strain behaves similarly to wild type animals.

For the fluorescence quantification, a Kruskal-Wallis test was performed to identify statistically significant differences (with a Benjamini, Krieger, Yekutieli two-stage set-up method to correct for multiple comparisons).

## Acknowledgements

We acknowledge Dr. Torsten Wittman and his lab for allowing Evangelie Chien to collect and analyze data using the spinning disc confocal scope. We thank L’Etoile lab members for their valuable comments and discussions.

---

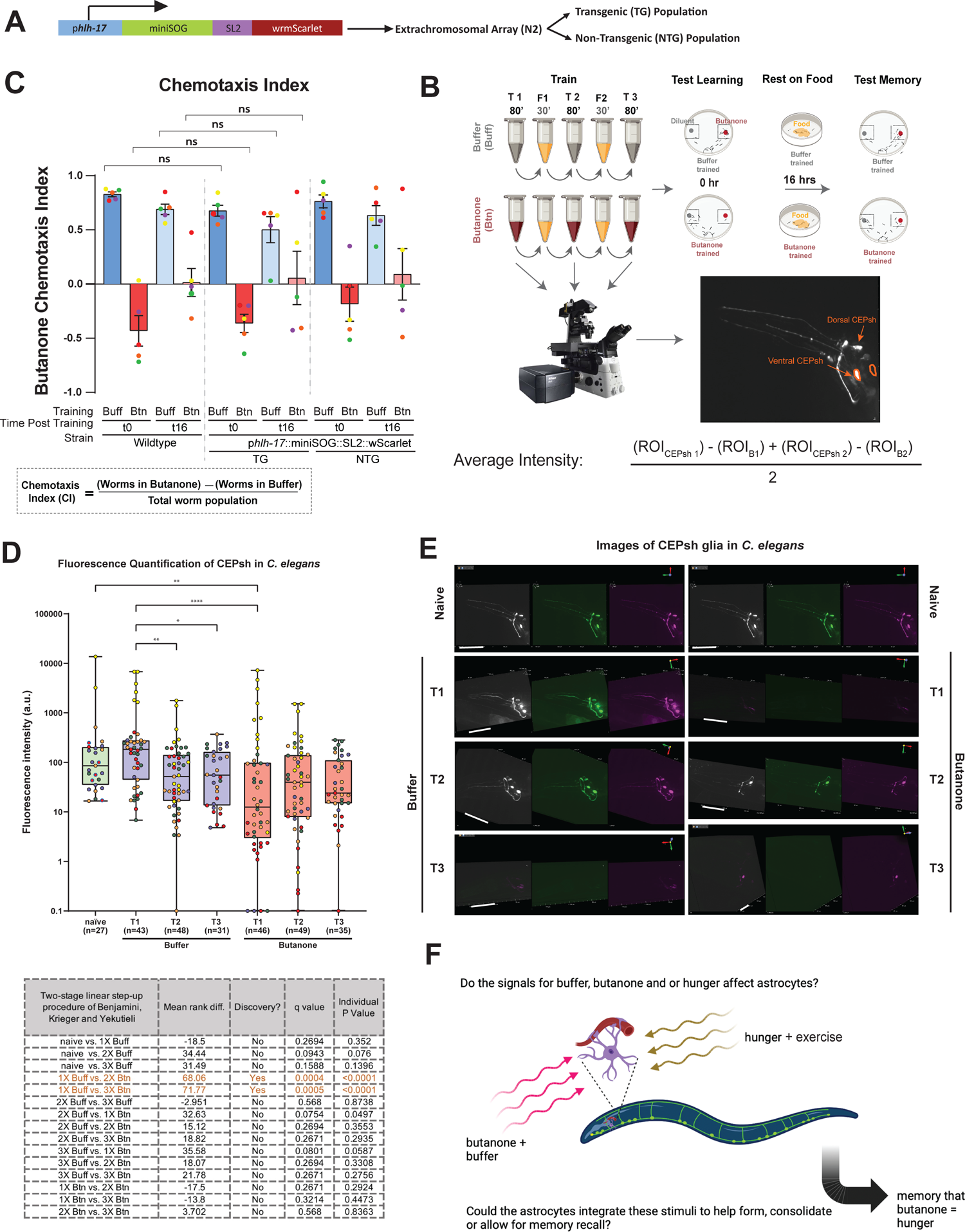

